# FlexAutoDock: A Flexible Platform for Automated Molecular Docking and Virtual Screening of Natural and Synthetic Compounds

**DOI:** 10.64898/2026.08.11.744098

**Authors:** Md. Feroj Ahmed, Md. Fahim Faysal, Khalid Muntasir Sawad, Md. Ashik -E- Elahi, Tasfia Noor, Md. Kaderi Kibria, Md. Mehedi Hasan, Md. Nurul Haque Mollah

## Abstract

Drug discovery (DD) is a complex, time-consuming, and resource-intensive process that involves the identification of therapeutic targets, selection of bioactive compounds, and extensive experimental validation. The discovery of promising therapeutic compounds from large libraries of phytochemicals and synthetic molecules remains a major challenge in modern drug development. Screening millions of compounds through conventional experimental approaches requires substantial time, cost, and computational resources. In recent years, in-silico molecular docking has emerged as an important computational approach for predicting interactions between small molecules and target proteins, thereby helping researchers prioritize promising compounds for further investigation. Several molecular docking webservers, including iScreen, SwissDock, CB-Dock2, DockThor, and MTiOpenScreen, have been developed to support virtual screening studies. However, many currently available platforms still face some important limitations. Most existing tools lack integrated repositories of medicinal plant-derived phytochemicals and organism-derived bioactive compounds, automated mapping between plants and their associated phytochemicals, and flexible ligand retrieval using chemical names, SMILES strings, PubChem CIDs, or drug names. In addition, many platforms require extensive manual protein and ligand preparation, provide limited support for AlphaFold-predicted protein structures, and lack efficient large-scale multi-target virtual screening. Most existing docking platforms offer limited support for interactive inspection of docked protein-ligand complexes, often requiring users to download the results and analyse them using external molecular visualization software. To address these limitations, we developed FlexAutoDock, an automated cloud-based molecular docking platform that provides a unified environment for protein-ligand docking and large-scale virtual screening. Unlike existing web servers, FlexAutoDock integrates curated repositories of medicinal plant-derived phytochemicals, organism-derived bioactive compounds, and synthetic compounds from the ZINC database while supporting flexible ligand acquisition through medicinal plant or organism selection, chemical names, SMILES strings, PubChem CIDs, and drug-name queries. The platform further streamlines the docking workflow through automated protein structure retrieval from the Protein Data Bank and AlphaFold databases, receptor and ligand preparation, chain-specific protein selection, blind and site-specific docking, interactive visualization of predicted protein-ligand complexes, and scalable multi-target virtual screening. The resulting platform enables rapid, flexible, and large-scale virtual screening while simplifying the molecular docking workflow, providing researchers with an accessible computational resource for accelerating early-stage drug discovery. FlexAutoDock offers a fast, reliable, and accessible computational platform for molecular docking and virtual screening, freely available to the scientific community at http://103.99.177.82:3000/.

## Introduction

Molecular docking (MD) is a computational prediction method widely used to investigate the interaction between small molecules (e.g., ligands) and target proteins. The primary objective of molecular docking is to predict the most favourable binding orientation and conformation of a ligand within the active site of a receptor protein, enabling the formation of a stable protein-ligand complex and providing insights into molecular-level interactions[1]. MD plays a crucial role in structure-based DD by helping researchers to prioritize the most promising therapeutic compounds from millions of candidate molecules for further experimental validation[2]. This approach significantly reduces the time, cost, and experimental effort required during the DD process[2,3]. Along with synthetic and inorganic chemical compounds, natural products derived from medicinal plants also represent an important source of DD. Medicinal plants distributed across diverse geographical regions contain a vast array of structurally diverse bioactive phytochemicals with significant pharmacological potential[4]. Traditionally, different populations worldwide have used various medicinal plants to treat similar or distinct diseases, suggesting the presence of biologically active compounds with therapeutic relevance[5,6]. Consequently, medicinal plant-derived phytochemicals represent valuable resources for the identification and development of novel drug candidates[7]. Nevertheless, identifying the most promising drug candidates from millions of natural and synthetic compounds remains a difficult and time-consuming task. Researchers often need to manually collect plant-specific phytochemicals and other chemical compounds from multiple databases and subsequently preprocess them into docking-compatible formats before MD analysis can be conducted[3,8]. In addition, preparing disease-associated target proteins for MD is another challenging and labour-intensive procedure, especially when handling large numbers of proteins simultaneously. Protein preparation generally requires several preprocessing steps, including removal of unnecessary atoms and water molecules, chain selection, charge assignment, active-site identification, and grid-box generation to ensure the proteins are suitable for docking analysis[9,10].

To address these challenges, researchers and bioinformaticians have developed various MD and virtual screening webservers and software platforms. Among the most widely used tools SwissDock[11], CB-Dock2[12], DockThor[13], MTiOpenScreen[14], and iScreen[15], are most famous types of tools and software. And these platforms have become important components of modern structure-based DD workflows by enabling researchers to prioritize promising compounds before proceeding to costly and time-intensive experimental validation studies. Despite substantial progress in molecular docking technologies, most currently available virtual screening and docking platforms still lack integrated global medicinal plant-phytochemical repositories and automated plant-to-compound mapping systems [11–15]. Consequently, researchers are often required to manually collect, curate, and preprocess phytochemical datasets from multiple fragmented databases before docking analysis can be performed. This limitation not only increases the complexity, computational burden, and time required for large-scale phytochemical screening, but also restricts the systematic exploration of region-specific medicinal plants and their associated bioactive compounds. As a result, the efficient translation of ethnopharmacological knowledge into scalable computational drug discovery remains a major unresolved challenge in current virtual screening workflows.

Additionally, this problem none of these platforms fully able to perform docking with multiple protein and multiple drugs docking[11–15]. Following these major concerns all tools have distinct type of disadvantages like some tools cannot curated multiple drug’s structure automatically[11,12], though some of these have integrated limited number of local repositories[13,15].

To overcome these limitations, we developed FlexAutoDock, an automated cloud-based molecular docking platform that provides a unified framework for protein-ligand docking and large-scale virtual screening. The platform integrates curated repositories of medicinal plant-derived phytochemicals, organism-derived bioactive compounds, and synthetic compounds from the ZINC database, while supporting flexible ligand acquisition through medicinal plant selection, chemical names, SMILES strings, PubChem CIDs, and drug-name queries. FlexAutoDock further streamlines the docking workflow by automating protein structure retrieval from the Protein Data Bank (PDB) and AlphaFold databases, protein and ligand preparation, chain-specific protein selection, blind and site-specific docking, and scalable multi-target virtual screening. By integrating these capabilities into a single platform, FlexAutoDock minimizes manual intervention, improves the efficiency and reproducibility of virtual screening, and provides a comprehensive computational resource for natural product research, drug repurposing, and structure-based DD.

## Materials and Methods

### Global Natural Compound databases

To facilitate the global exploration of medicinal plants, we have curated comprehensive information on plant species and their associated bioactive compounds from multiple publicly available databases. These data sources, which originate from different geographical regions, have been systematically integrated to create a large repository of plant-phytochemical information for research and educational purposes. The repository now contains extensive plant-specific chemical data collected from diverse regions of the world, providing researchers with a centralized platform for exploring medicinal plants and their phytoconstituents. This resource enables users to investigate which plant-derived compounds may exhibit therapeutic potential against particular diseases. The total curated data information given below (**see Table 1**).

**Table 1.** Databases curated to create global plant database.

| Database | Number of plants/organisms | Number of compounds | Regions |
| --- | --- | --- | --- |
| COCONUT[16] | 71448 | 738827 | Worldwide |
| IMPPAT[17] | 4010 | 17967 | India |
| NPASS [18] | 48311 | 203386 | Worldwide |
| TCMSP[19] | 498 | 6477 | China |
| CMAUP[20] | 6807 | 2979 | China |
| VIETHERB[21] | 1238 | 10262 | Vietnam |
| TPPT[22] | 1565 | 1565 | Switzerland and Central Europe |

### Inputs

The primary objective of the FlexAutoDock webserver is to reduce the time, cost, and labour associated with the DD process by providing an automated platform for molecular docking of multiple proteins and ligands. The webserver simplifies the complex computational procedures involved in docking studies, making them more accessible and user-friendly. As a result, researchers from diverse academic backgrounds, including those with limited computational expertise, can easily perform docking analyses without extensive technical knowledge. Furthermore, the FlexAutoDock webserver offers versatile input options, allowing users to upload multiple protein structures and ligand molecules in various formats. This flexibility enables researchers to conduct large-scale screening studies efficiently while minimizing the technical barriers commonly associated with computational DD workflows.

### Input of Natural Products as Drug Candidates

For plant-based docking, users simply need to select the scientific name of a medicinal plant. If the selected plant is available in the curated global plant–chemical database, the webserver displays a list of unique bioactive compounds associated with that plant. Users can either select all available compounds simultaneously or choose specific molecules of interest according to their research objectives. In addition, the platform allows users to explore phytochemicals from multiple medicinal plants within a single workflow. This feature enables researchers to compare compounds from different plant sources and identify which plant-derived chemicals exhibit stronger binding affinities toward a target protein. By providing easy access to diverse phytochemicals and flexible compound selection options, the webserver facilitates efficient plant-based virtual screening and accelerates the identification of promising natural drug candidates.

### Input of Chemical Names as Drug Candidates

Users can also search and select drug candidates directly by entering the chemical name of a compound. The webserver initially searches the queried compound within the curated global plant–chemical database. If the compound is found, the corresponding information is retrieved and made available for docking analysis. In cases where the requested compound is not present in the internal database, the system automatically searches the PubChem database through its REST API to obtain the relevant chemical information and molecular structure. This integrated search strategy ensures that users can access a wide range of compounds, including both plant-derived molecules and externally available chemical entities. By combining the curated plant– chemical repository with real-time retrieval from external databases, the webserver provides a flexible and comprehensive platform for selecting potential drug candidates for molecular docking studies.

### Input of PubChem CID as Drug Candidates

The webserver also allows users to retrieve and submit candidate drug molecules using PubChem Compound Identifiers (CIDs). Users may enter individual CID numbers manually or upload multiple CIDs simultaneously in text or CSV format for batch processing. Upon submission, the system automatically retrieves the necessary chemical information and molecular structures from the PubChem database through its REST API. The downloaded compounds are then processed and prepared for molecular docking without requiring additional user intervention. This automated workflow enables efficient large-scale screening of compounds and significantly reduces the time and effort involved in ligand preparation.

### Input of ZINC IDs as Drug Candidates

The webserver also enables users to retrieve and submit candidate drug molecules using ZINC IDs. Since the platform is integrated with the ZINC database, users can enter individual ZINC IDs manually or upload multiple ZINC IDs simultaneously in text or CSV format for batch processing. Upon submission, the system automatically retrieves the corresponding SMILES representations and other necessary chemical information from the ZINC database. The compounds are then processed and prepared for molecular docking without requiring additional user intervention. This automated workflow facilitates high-throughput virtual screening and substantially reduces the time and effort associated with ligand preparation.

### Input of SMILES as Drug Candidates

The FlexAutoDock webserver also incorporates a dedicated pipeline for processing SMILES representations of chemical compounds for molecular docking studies. Users can either enter individual SMILES strings manually or upload multiple SMILES in batch mode using a text file. Upon submission, the system automatically converts the SMILES strings into appropriate three-dimensional molecular structures and performs the necessary ligand preparation steps required for docking analysis. This automated process eliminates the need for additional software or manual preprocessing, making the workflow more efficient and user-friendly.

A visual overview of the multiple ways of drug candidate selection interface on the FlexAutoDock server given below (**see Figure 2**).

**Figure 1.**
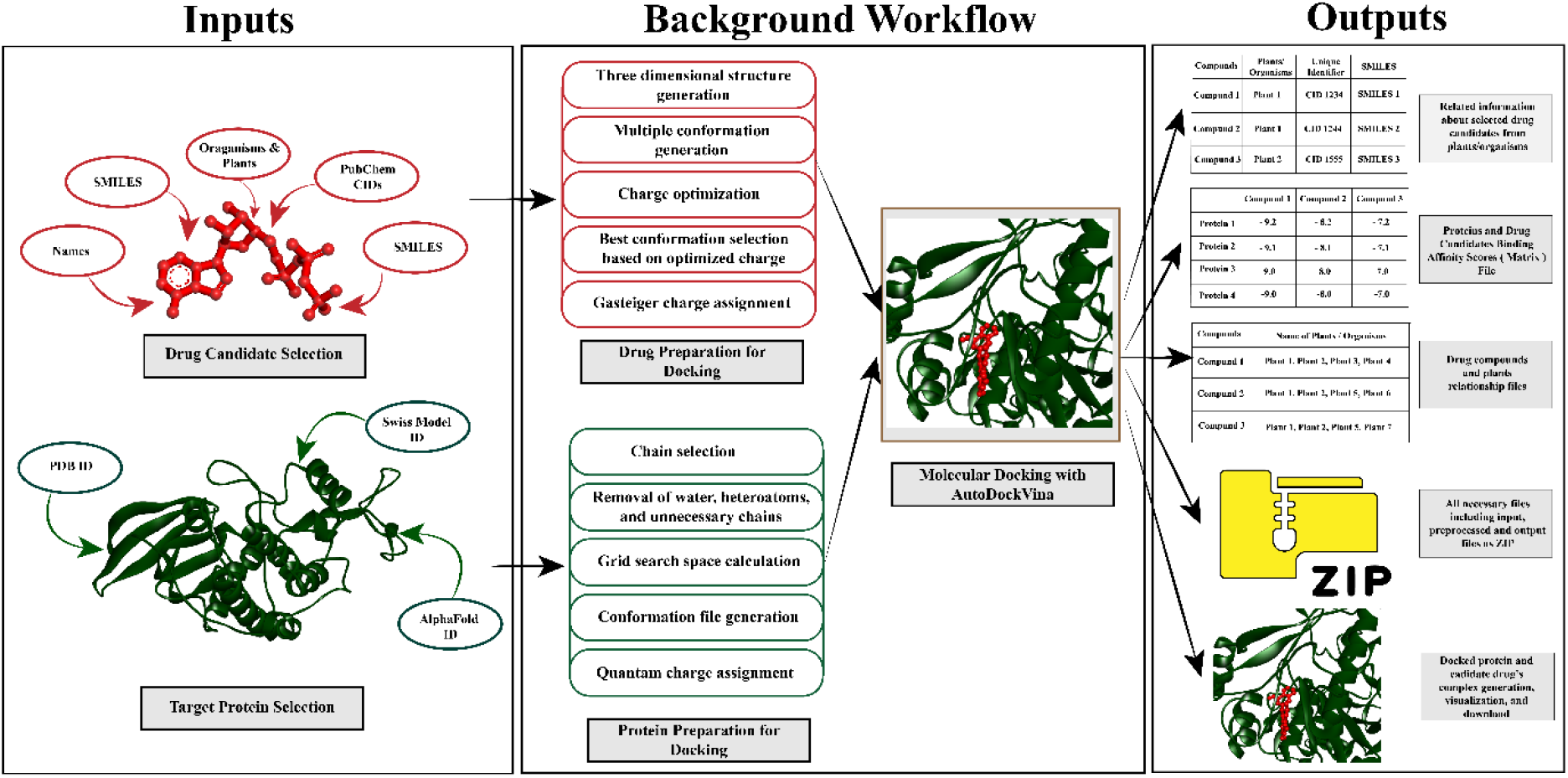
Schema describing the features of the FlexAutoDock webtool including inputs, background workflow, and outputs.

**Figure 2.**
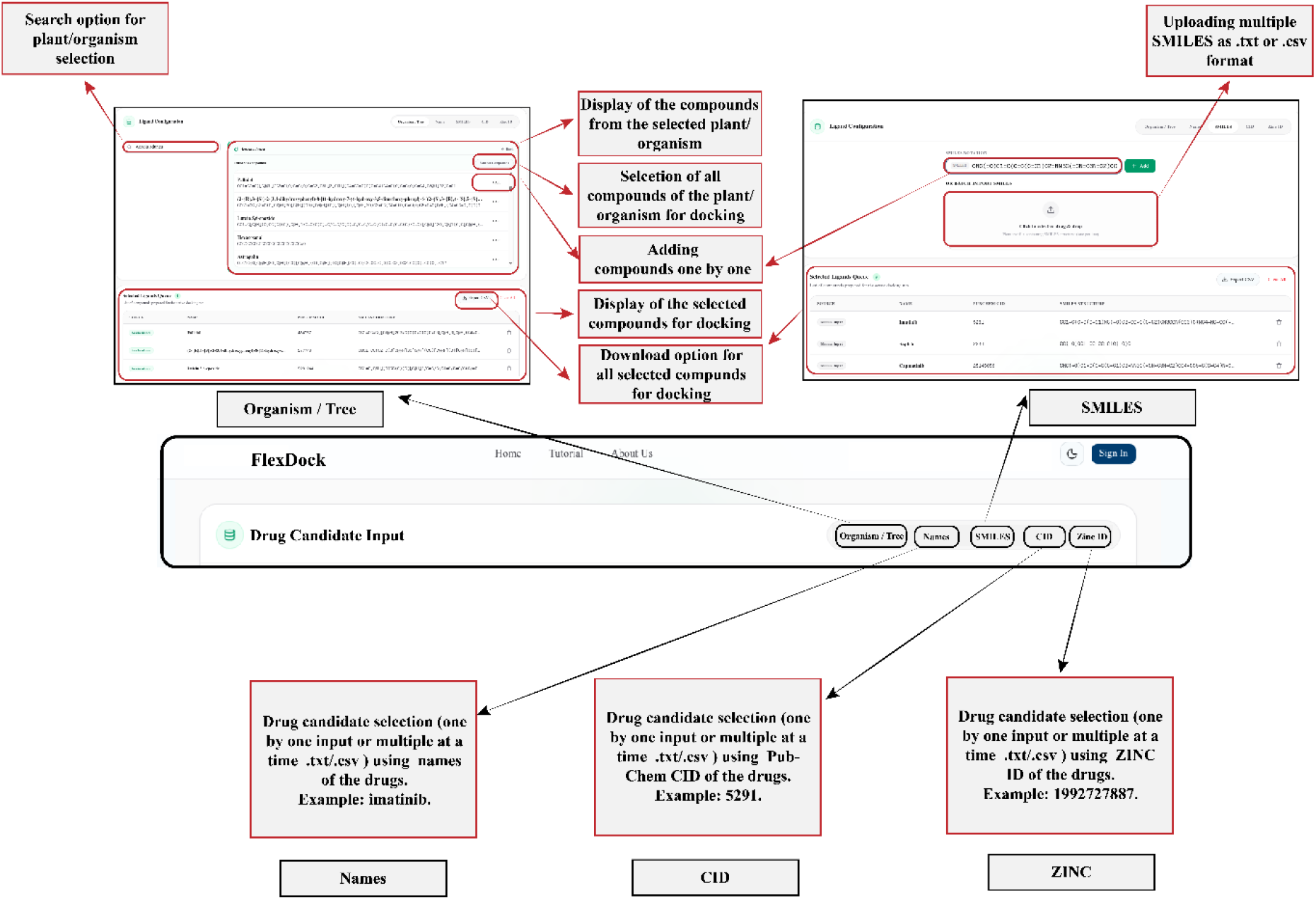
Overview of the multiple ways of drug candidate selection interface on the FlexAutoDock server.

### Input of Proteins as Drug Targets

The FlexAutoDock webserver provides substantial flexibility for the selection and preparation of protein targets for molecular docking studies. Users can easily submit one or multiple proteins as potential drug targets by simply providing the corresponding PDB ID or AlphaFold ID of the protein structure. Upon submission, the platform automatically retrieves the required protein structures from the Protein Data Bank (PDB) and AlphaFold databases through their respective REST APIs. After the structures are obtained, users can select the specific protein chain to be used for docking analysis. The webserver supports both blind docking and active-site docking approaches. For active-site docking, users can specify the center coordinates (x, y, and z) along with the grid box dimensions to define the binding region of interest. Alternatively, users may choose blind docking, in which the system automatically generates an appropriate search space covering the entire protein structure. To assist users in defining the docking region, the platform also provides a visualization option that allows the selected search space or active-site coordinates to be inspected before docking. Following these selections, all protein structures are automatically processed and prepared for molecular docking, eliminating the need for manual intervention and simplifying the overall docking workflow. A visual overview of the target protein submission and preprocessing configuration interface on the FlexAutoDock server showed below (**see Figure 3**).

**Figure 3.**
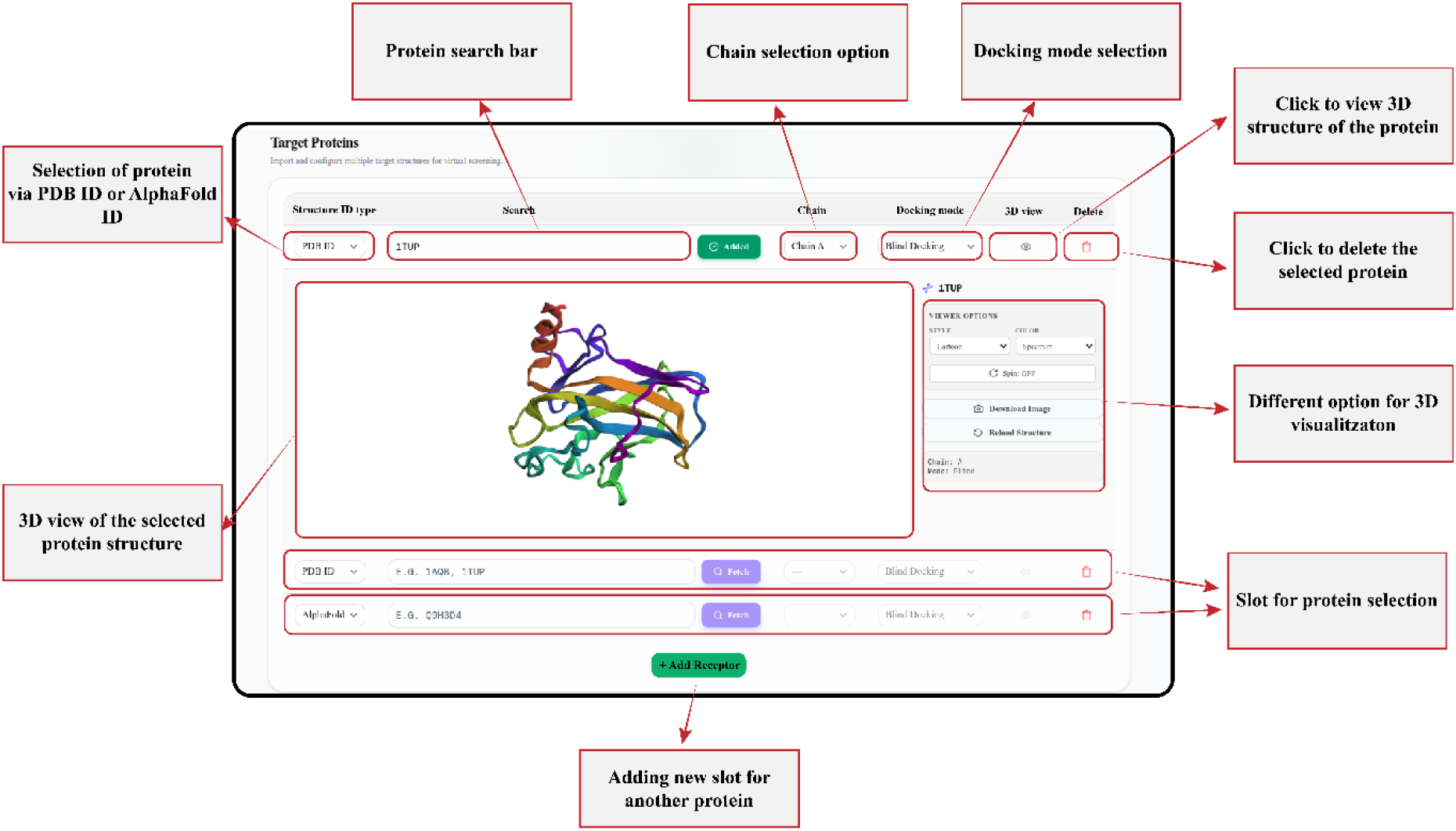
Overview of the target protein submission and preprocessing configuration interface on the FlexAutoDock server.

## Structure Preparations for Docking

### Protein Preparation

After the user submission, the FlexAutoDock webserver automatically prepares the target protein for molecular docking studies. This automated preprocessing step eliminates the need for manual protein preparation and ensures that the structures are suitable for docking analysis. Initially, the system retains only the user-selected protein chain and removes all unwanted chains from the structure. In addition, water molecules, heteroatoms, and other non-essential components are automatically excluded to minimize potential interference during the docking process and to generate a clean receptor structure. Subsequent preprocessing steps are performed using publicly available functions from AutoDockTools[23]. These procedures include the addition of polar hydrogen atoms, assignment of Kollman charges, and the generation of receptor PDBQT files required for molecular docking. By integrating these standardized preparation steps into an automated pipeline, the FlexAutoDock webserver provides a reliable, reproducible, and user-friendly framework for protein preparation without requiring any manual intervention.

### Drug Molecule Preparation

After user submission, the three-dimensional (3D) structures of drug molecules are either retrieved directly from the PubChem database or generated from SMILES representations using the RDKit Python library[24]. This flexible approach enables the platform to process compounds obtained from multiple sources while maintaining a standardized ligand preparation workflow. Following the generation of the initial 3D structures, hydrogen atoms are added to the molecules to ensure proper chemical representation. Subsequently, multiple molecular conformations (20 conformers) are generated and subjected to geometry optimization to identify energetically favourable structures. The optimized conformers are then evaluated, and the lowest-energy conformation is selected as the representative structure for docking analysis. Finally, the ligand structure is converted from PDB format to the PDBQT format required for molecular docking. During this conversion, Gasteiger charges are assigned using Open Babel[25], thereby preparing the ligand for subsequent docking studies. This automated ligand preparation pipeline ensures consistent and reliable processing of drug candidates while minimizing the need for manual intervention and reducing the complexity of molecular docking workflows.

### Molecular Docking

Molecular docking between the target proteins and candidate drug compounds is performed using AutoDock Vina[26]. The FlexAutoDock webserver allows users to customize the docking process by selecting the desired exhaustiveness value during job submission, thereby providing flexibility in balancing docking accuracy and computational time. The docking configuration file is generated automatically during the protein preparation stage. For blind docking, the system automatically determines the search space and creates the corresponding configuration file to cover the entire protein surface. In contrast, for active-site-specific docking, the configuration file is generated according to the user-provided coordinates and grid box dimensions, enabling focused docking within a defined binding region. After generating the appropriate configuration parameters, AutoDock Vina performs the docking simulations and predicts the binding conformations and binding affinities of the candidate compounds against the selected target proteins. This automated workflow simplifies the docking process while ensuring flexibility for both exploratory blind docking and targeted active-site docking studies. This docking results are completely reproducible and with the same input all the result will be same if used one to reproduce the results.

### Validation of the Docking Protocols

The FlexAutoDock web server employs AutoDock Vina[26] as its molecular docking engine. Since the underlying docking algorithm of this software has been extensively validated and widely adopted in computational DD, so this present validation was designed to assess the reliability of the FlexAutoDock implementation and its automated workflow rather than to revalidate the AutoDock Vina algorithm itself. To evaluate the performance of the docking workflow, 10 experimentally determined protein-ligand complexes with co-crystallized ligands were selected from the PDB (**see Table 2**). These complexes represented diverse protein families and binding-site architectures to assess the robustness and general applicability of the platform. For each complex, the native ligand was extracted from the crystal structure with PyMOL[27], while the corresponding receptor was prepared using the automated protein preparation pipeline implemented in FlexAutoDock. The binding site for each receptor was identified based on the coordinates of the co-crystallized ligand using BIOVIA Discovery Studio Visualizer[28], and the resulting centre coordinates (X, Y, and Z) were used to define the docking search space (**see Table 3**). Molecular docking was subsequently performed using AutoDock Vina with the search box cantered on the experimentally determined ligand-binding site. For each protein-ligand complex, the top 10 docking poses ranked by binding affinity were generated. The predicted poses were individually superimposed onto the corresponding crystallographic ligand, and the root mean square deviation (RMSD) between the heavy atoms of the docked and native ligand conformations was calculated using PyMOL. The docking pose exhibiting the lowest RMSD was considered the representative pose for validation. An RMSD value below 2.0 Å was regarded as an excellent reproduction of the experimental binding mode, whereas RMSD values below 3.0 Å were considered acceptable for validating the docking workflow[29]. The docking protocol was considered successful when the experimentally observed ligand binding orientation was accurately reproduced.

**Table 2.**
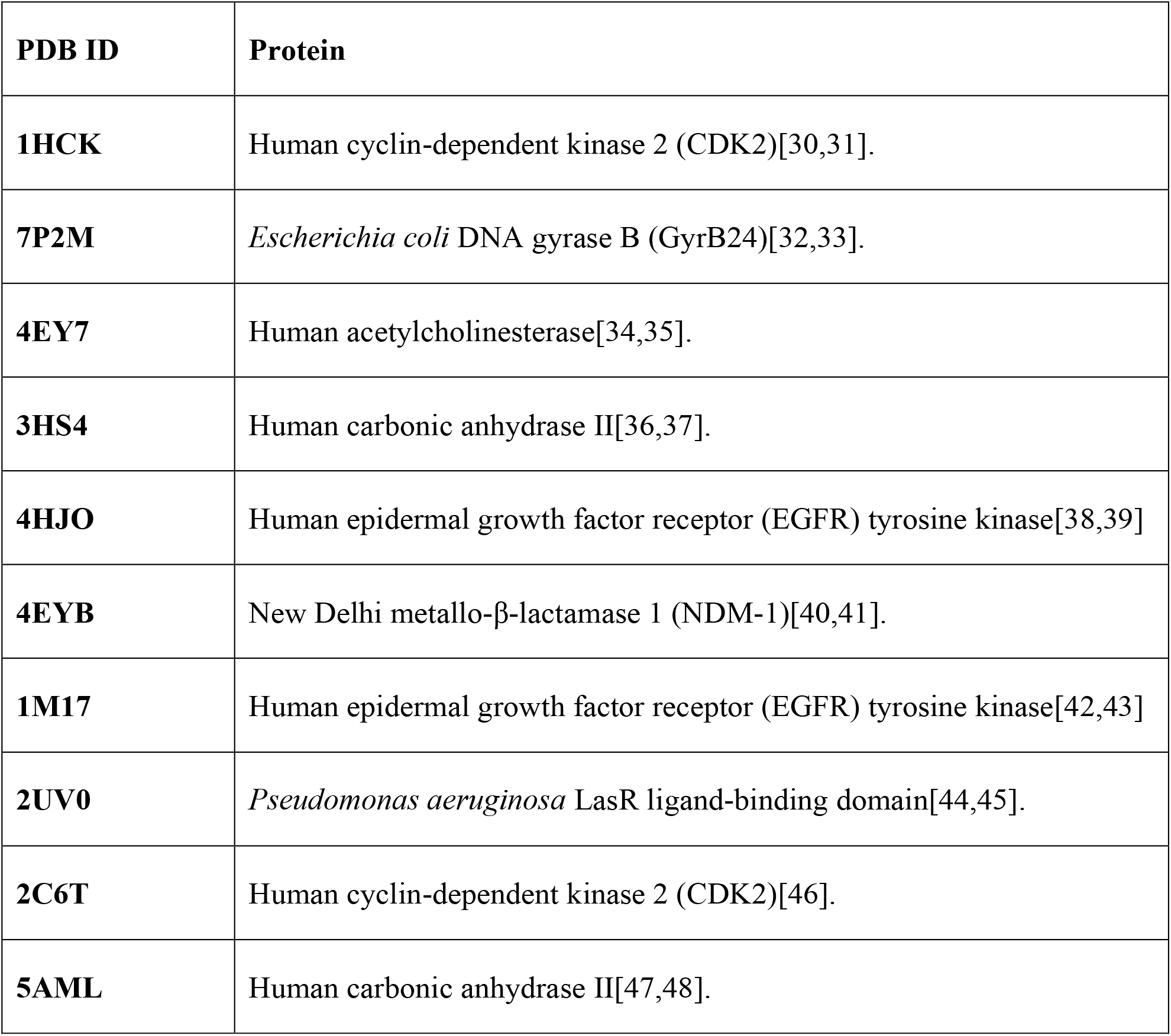
Experimentally determined protein structures used for docking protocol validation.

| **PDB ID** | **Protein** |
| --- | --- |
| **1HCK** | Human cyclin-dependent kinase 2 (CDK2)[30,31]. |
| **7P2M** | *Escherichia coli* DNA gyrase B (GyrB24)[32,33]. |
| **4EY7** | Human acetylcholinesterase[34,35]. |
| **3HS4** | Human carbonic anhydrase II[36,37]. |
| **4HJO** | Human epidermal growth factor receptor (EGFR) tyrosine kinase[38,39] |
| **4EYB** | New Delhi metallo-β-lactamase 1 (NDM-1)[40,41]. |
| **1M17** | Human epidermal growth factor receptor (EGFR) tyrosine kinase[42,43] |
| **2UV0** | *Pseudomonas aeruginosa* LasR ligand-binding domain[44,45]. |
| **2C6T** | Human cyclin-dependent kinase 2 (CDK2)[46]. |
| **5AML** | Human carbonic anhydrase II[47,48]. |

**Table 3.** Native ligand binding site coordination in the protein structure.

| **PDB ID** | **X Coord.** | **Y Coord.** | **Z Coord.** | **Radius (Å)** |
| --- | --- | --- | --- | --- |
| 1HCK | 100.557 | 97.7911 | 81.7647 | 7.645 |
| 7P2M | -18.6026 | -15.0457 | 6.55323 | 9.892 |
| 4EY7 | -10.6741 | -42.2676 | 30.5038 | 12.2 |
| 3HS4 | -4.62293 | 1.60327 | 10.6063 | 7.3 |
| 4EYB | -2.44815 | 8.60857 | 23.5789 | 10.4 |
| 1M17 | 22.0137 | 0.25283 | 52.794 | 9.596 |
| 2UV0 | 21.5953 | 13.9975 | 82.7882 | 11.1 |
| 2C6T | 37.3169 | 134.101 | 32.1855 | 7.509 |
| 5AML | -4.71109 | 3.79718 | 14.3039 | 7.416 |
| 4HJO | 24.7702 | 9.19393 | -0.00331 | 9.497 |

## Results and Outputs

### Outputs

FlexAutoDock webserver provides a wide range outputs to the users including those outputs which are really necessary for further analysis in the DD pipeline. The outputs are listed below as.

### Compound Name Visualization and Download

By selecting one or more plant species in the FlexAutoDock server, the platform automatically retrieves and displays the associated compounds available for the selected species in the curated database. Users can download the displayed compound information by selecting the Save option before moving into the docking protocols.

### Binding Affinity Matrix (BAS) Visualization and Download

The FlexAutoDock server displays the BAS score of interaction between drug candidates and receptors proteins. Users can easily able to download the BAS matrix file by clicking the save option given above the file.

### Protein-Ligand Complex Visualization and Download

The FlexAutoDock server enables users to visualize the predicted binding pose of each docked ligand within the corresponding protein structure by clicking the associated Binding Affinity Score (BAS) in the BAS matrix. The resulting protein–ligand complex is displayed through an interactive molecular viewer, facilitating inspection of the predicted binding orientation. Furthermore, the docked protein-ligand complex can be downloaded in PDB format for subsequent structural analysis and post-docking investigations using external molecular visualization and computational tools.

### Zip Files Download

Upon completion of the docking process, users can download all input metadata and docking results as a compressed ZIP archive. The downloaded package contains the original protein structures (PDB), prepared protein structures (PDBQT), AutoDock Vina configuration files, ligand structures (PDB), prepared ligand structures (PDBQT), docking output files containing the top 10 ranked binding conformations for each ligand in PDBQT format, predicted binding affinity scores (BAS) for each protein–ligand pair, a comprehensive binding affinity score matrix summarizing all docking results, and a phytochemical-plant association file that links each screened phytochemical to its corresponding plant source.

A visual representaion of the interactive docking results and data export interface of the FlexAutoDock server (**see Figure 4**).

**Figure 4.**
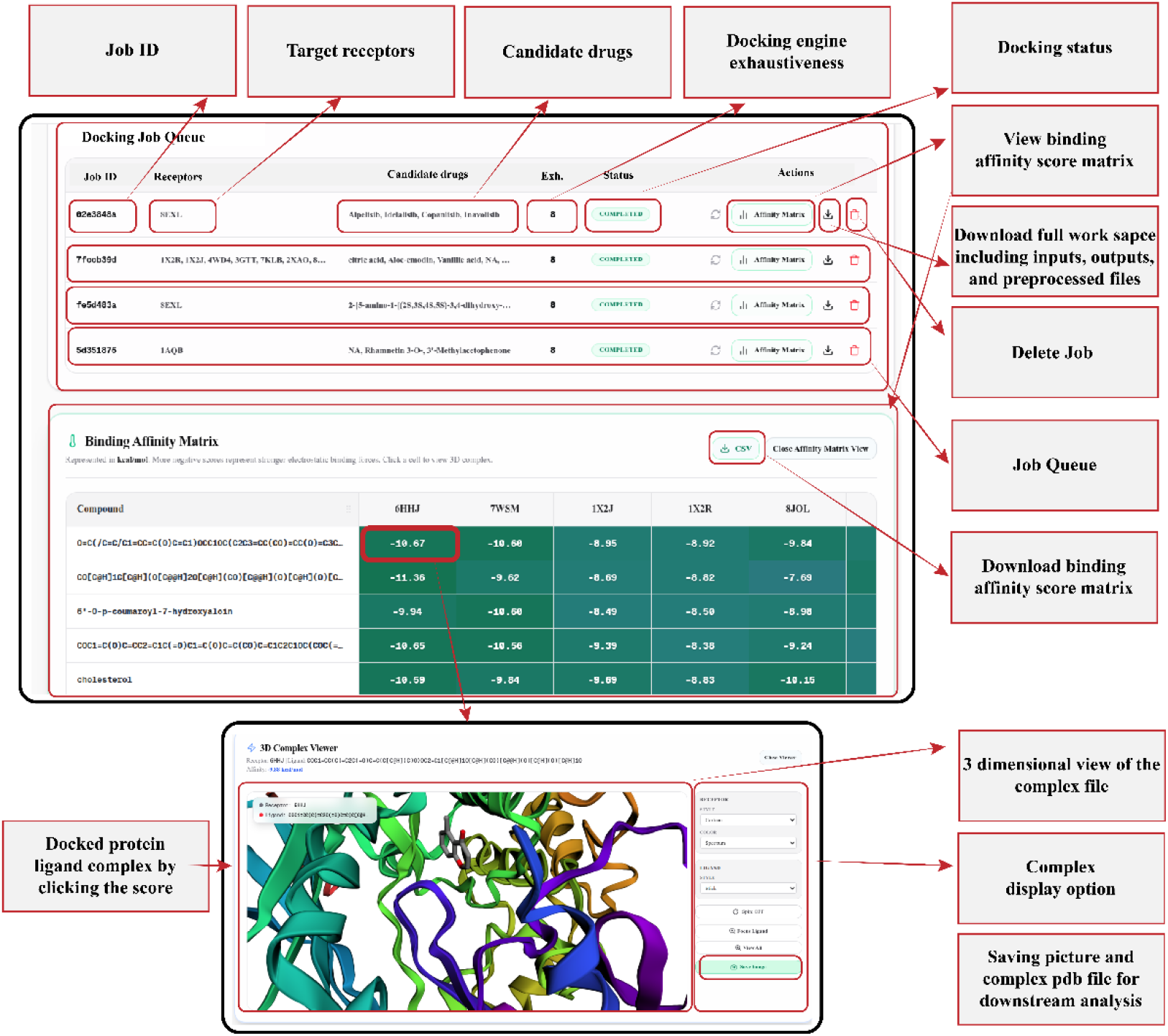
Overview of the interactive docking results and data export interface of the FlexAutoDock server.

## Validation of the Docking Protocol

The docking workflow implemented in FlexAutoDock was evaluated using ten experimentally resolved protein-ligand complexes obtained from the PDB. For each complex, the co-crystallized ligand was redocked into its corresponding binding pocket, and the generated docking poses were compared with the experimentally determined ligand conformation. Among the top 10 predicted binding poses generated for each complex, the pose exhibiting the lowest RMSD relative to the crystallographic ligand was selected for validation. The validation results (**see Table 4**) showed that the FlexAutoDock workflow consistently reproduced the experimentally observed ligand-binding orientations with high accuracy. The RMSD values ranged from 0.432 to 1.506 Å (**see Figure 5**), with all ten protein-ligand complexes achieving RMSD values below 2.0 Å, indicating accurate reproducibility of the crystallographic binding mode. The smallest deviation was obtained for PDB ID 4EY7 (0.432 Å), whereas the largest RMSD was observed for PDB ID 4HJO (1.506 Å). Five protein-ligand complexes (1HCK, 7P2M, 4EY7, 3HS4, and 4EYB) were accurately reproduced by the highest-ranked docking pose (Pose 1), whereas the remaining five complexes (1M17, 2UV0, 2C6T, 5AML, and 4HJO) were best represented by Pose 3 or Pose 4. The predicted BAS ranged from -7.0 to -12.2 kcal/mol, demonstrating energetically favourable binding conformations across all validation complexes. Notably, the low RMSD values obtained for structurally diverse proteins, indicate that the FlexAutoDock workflow can accurately reproduce experimentally determined binding modes across a broad range of protein families and binding-site architectures. These results confirm the reliability and robustness of the automated docking workflow implemented in the FlexAutoDock platform.

**Figure 5.**
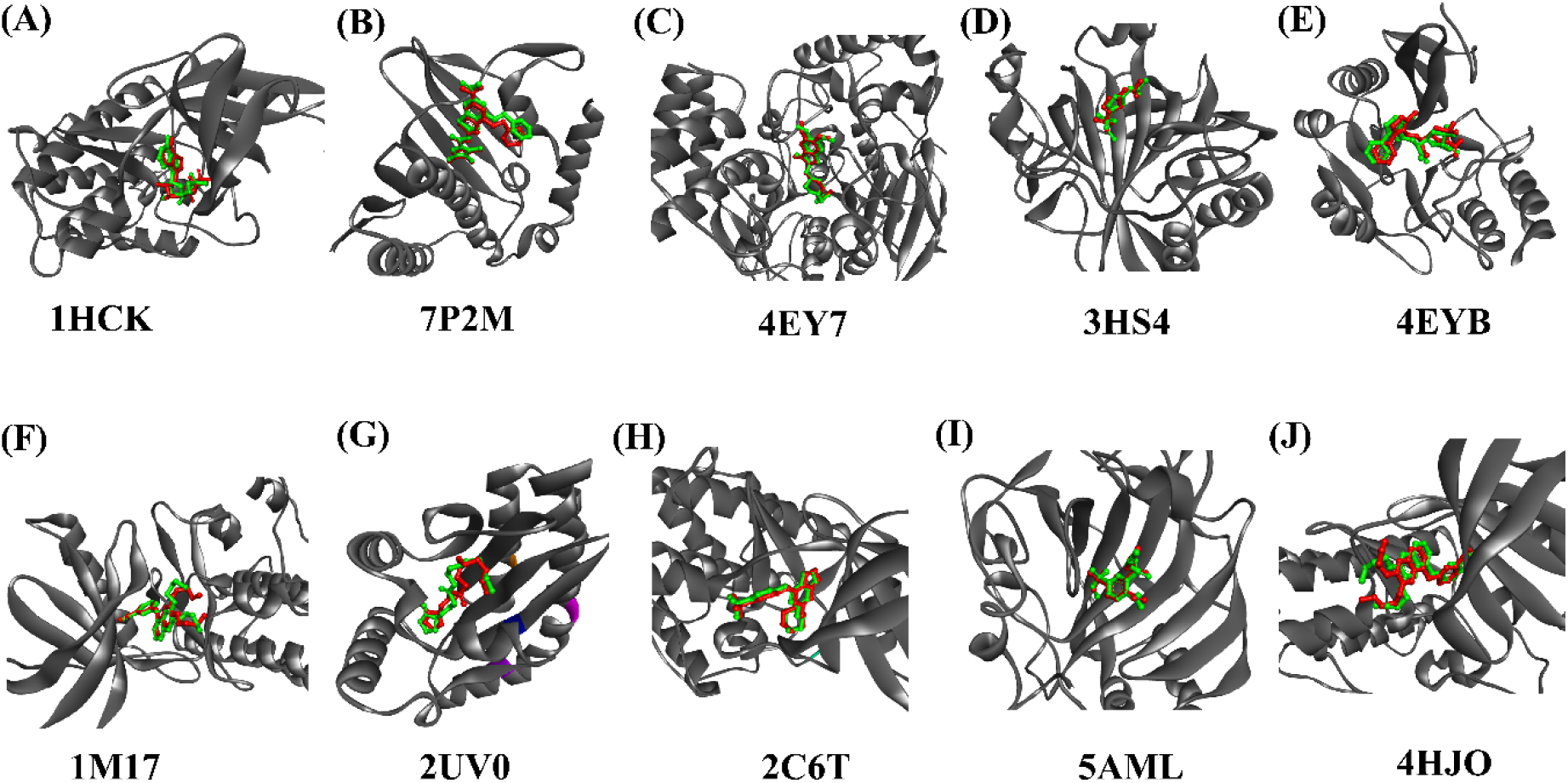
Three-dimensional view of docked ligands superimposed against reference co-crystallized structures for proteins (A) HCK, (B) 7P2M, (C) 4EY7, (D) 3HS4, (E) 4EYB, (F) 1M17, (G) 2UV0, (H) 2C6T, (I) 5AML, and (J) 4HJO.

**Table 4.** RMSD value between native ligand and redocked ligand for docking protocol validation.

| PDB ID | POSE | RMSD | BAS<br>(kcal/mol) |
| --- | --- | --- | --- |
| 1HCK | POSE 1 | 1.223 (31 to 31 atoms) | -8.6 |
| 7P2M | POSE 1 | 1.311 (31 to 31 atoms) | -9 |
| 4EY7 | POSE 1 | 0.432 (28 to 28 atoms) | -12.2 |
| 3HS4 | POSE 1 | 1.409 (13 to 13 atoms) | -7.5 |
| 4EYB | POSE 1 | 1.323 (29 to 29 atoms) | -8 |
| 1M17 | POSE 3 | 1.236 (29 to 29 atoms) | -7 |
| 2UV0 | POSE 3 | 1.264 (21 to 21 atoms) | -8.6 |
| 2C6T | POSE 4 | 0.859 (27 to 27 atoms) | -8.5 |
| 5AML | POSE 4 | 1.201 (21 to 21 atoms) | -7.50 |
| 4HJO | POSE 4 | 1.506 (29 to 29 atoms) | -7.5 |

## Conclusion & Discussion

FlexAutoDock is a freely accessible and user-friendly web server developed to facilitate molecular docking and large-scale virtual screening for computational DD. The platform integrates locally curated repositories of phytochemicals derived from medicinal plants, bioactive compounds derived from diverse organisms, and synthetic compounds obtained from the ZINC database, while also supporting user-defined ligands through chemical names, SMILES strings, and PubChem CIDs. By integrating automated protein structure retrieval, receptor and ligand preparation, binding-site definition, and molecular docking within a unified workflow, FlexAutoDock substantially reduces the technical expertise, computational effort, and time required to perform virtual screening most promising drug candidates. The FlexAutoDock utilizes the most used and reliable software autodock vina as it’s docking engine which eventually make this server reliable tool for virtual screening of large numbers of drug candidates for selecting the most promising drug candidates among them. In addition, the docking protocol implemented in FlexAutoDock was independently validated by redocking co-crystallized ligands into their corresponding experimentally determined protein structures. The redocked ligand poses closely reproduced the orientations observed in the crystal structures, demonstrating the reliability and accuracy of the docking protocol. This streamlined workflow enables researchers to efficiently prioritize promising candidate molecules for subsequent experimental validation, thereby supporting natural product research, drug repurposing, and structure-based DD. We anticipate that FlexAutoDock will serve as a valuable computational resource for the scientific community by facilitating efficient virtual screening and accelerating the identification of potential therapeutic candidates for future experimental and translational research.

## Authors contribution

**Conceptualization:** Md. Feroj Ahmed and Md Nurul Haque Mollah; **Data curation:** Md. Feroj Ahmed, Md. Fahim Faysal, Khalid Muntasir Sawad, and Md. Ashik -E- Elahi ; **Formal analysis:** Md. Feroj Ahmed and Md Nurul Haque Mollah ; **Investigation:** Md. Feroj Ahmed, Md. Fahim Faysal, Khalid Muntasir Sawad, Md. Ashik -E- Elahi, and Tasfia Noor; **Methodology**: Md. Feroj Ahmed, Khalid Muntasir Sawad, Md. Ashik -E- Elahi, and Md Nurul Haque Molla; **Project administration:** Md Nurul Haque Mollah; **Resources:** Md. Feroj Ahmed, and Md. Nurul Haque Mollah ; **Software:** Md. Feroj Ahmed, Md. Fahim Faysal, Khalid Muntasir Sawad, Md. Ashik -E- Elahi, and Tasfia Noor; **Supervision:** Md Nurul Haque Mollah; **Validation:** Md. Feroj Ahmed, Tasfia Noor, and Md Nurul Haque Mollah; **Visualization:** Md. Feroj Ahmed, Md. Fahim Faysal, Khalid Muntasir Sawad, and Md. Ashik -E- Elahi; **Writing – original draft:** Md. Feroj Ahmed, and Md Nurul Haque Mollah,; **Writing – review & editing:** Md. Feroj Ahmed, Md. Kaderi Kibria, Md. Mehedi Hasan, and Md Nurul Haque Mollah.

## Ethics statement

Not Applicable

## Consent to Publish

Not Applicable

